# Physiomarkers in Real-Time Physiological Data Streams Predict Adult Sepsis Onset Earlier Than Clinical Practice

**DOI:** 10.1101/322305

**Authors:** Franco van Wyk, Anahita Khojandi, Robert L. Davis, Rishikesan Kamaleswaran

## Abstract

**Rationale:** Sepsis is a life-threatening condition with high mortality rates and expensive treatment costs. To improve short- and long-term outcomes, it is critical to detect at-risk sepsis patients at an early stage.

**Objective:** Our primary goal was to develop machine learning models capable of predicting sepsis using streaming physiological data in real-time.

**Methods:** A dataset consisting of high-frequency physiological data from 1,161 critically ill patients admitted to the intensive care unit (ICU) was analyzed in this IRB-approved retrospective observational cohort study. Of that total, 634 patients were identified to have developed sepsis. In this paper, we define sepsis as meeting the Systemic Inflammatory Response Syndrome (SIRS) criteria in the presence of the suspicion of infection. In addition to the physiological data, we include white blood cell count (WBC) to develop a model that can signal the future occurrence of sepsis. A random forest classifier was trained to discriminate between sepsis and non-sepsis patients using a total of 108 features extracted from 2-hour moving time-windows. The models were trained on 80% of the patients and were tested on the remaining 20% of the patients, for two observational periods of lengths 3 and 6 hours.

**Results:** The models, respectively, resulted in F1 scores of 75% and 69% half-hour before sepsis onset and 79% and 76% ten minutes before sepsis onset. On average, the models were able to predict sepsis 210 minutes (3.5 hours) before the onset.

**Conclusions:** The use of robust machine learning algorithms, continuous streams of physiological data, and WBC, allows for early identification of at-risk patients in real-time with high accuracy.

## 1. Introduction

In the critical care environment, the availability of vast volumes of data present a unique opportunity to generate novel insights for better care [1, 2], The analysis of such substantial volumes of data are now more tractable with the use of sophisticated and efficient machine learning methods and strategies [1, 3–5]. The management of sepsis, one of the most expensive and life-threatening medical conditions, can benefit from the use of such tools, specifically to identify at risk patients earlier. If recognition is delayed, sepsis can rapidly progress to multiple organ dysfunction (MOD), which has a mortality rate as high as 80% [6, 7], An increase of approximately 8% in mortality rate is observed for each hour of delayed diagnosis of sepsis [8]. Predictive analytics applied to routinely collected data, such as physiological data, can reduce recognition gaps while allowing for targeted and early goal-directed therapy, while improving situational awareness in critical care.

Machine learning techniques have been extensively used in medical decision making and treatment planning. For instance, these algorithms have been used to predict at risk patients or patient outcomes, and to reduce alarm fatigue [9–11]. Similarly, machine learning algorithms have been successfully implemented in various medical image analyses to assist diagnosis and therapy in neurology, cardiology, and the detection of various cancers [12–19]. Machine learning algorithms have also shown promise in detecting/predicting sepsis [20, 21]. However, much of the recent work has been centered around static and often manually entered electronic health record (EHR) data [22].

Recent work has shown that ‘physiomarkers,’ such as reduced heart rate variability, may herald the onset of sepsis [23, 24], enabling a window of early recognition and treatment. In this paper, we utilize machine learning to identify early pathological characteristics in patients with sepsis, using continuous minute-by-minute data captured at the bedside. We define the onset of sepsis as meeting Systemic Inflammatory Response Syndrome (SIRS) criteria in the presence of the suspicion of infection, indicated by relevant ICD10 codes appearing in the primary and all additional discharge columns (see Supplemental Tables 1 and 2). We make the following key contributions in the area of precision medicine as applied to critical care medicine:

1. A predictive sepsis model built using a minimal set of continuous, routinely collected bedside physiological data streams
2. An analysis pipeline tailored for ‘online’ implementation
3. Identification of salient physiomarkers that predict the onset of sepsis in critically ill adults using an integrated machine learning approach

## 2. Materials and Methods

Figure 1 presents an overview of our approach. Physiological data including temperature, heart rate, systolic blood pressure (SBP), and respiratory rate, and laboratory data such as white blood cell (WBC) count are continuously used to make predictions about sepsis onset. Specifically, a fixed-width moving time-window of data is used as an input to the model, where the output is the probability of developing sepsis. Each minute, new observations in the form of physiological data are made on the patient at the bedside. Hence, the time window moves forward to include the data corresponding to these new observations, while the earliest data are removed from the time-window to maintain its width. Once these input data are updated, a new prediction about the likelihood of the patient developing sepsis is made.

**Figure 1.**
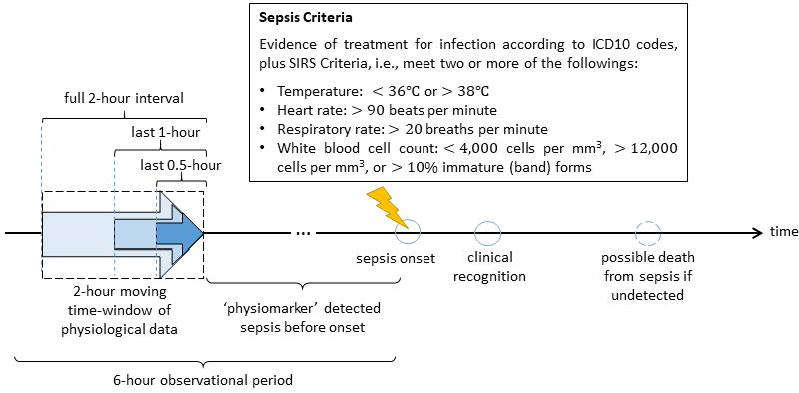
An overview of the approach (with a 6-hour observational period).

### 2.1 Data

Continuous minute-by-minute physiological data was captured using a proprietary Cerner CareAware iBus^®^ platform [25] at the Methodist LeBonheur Healthcare (MLH) System in Memphis TN between February and December 2017. The data was collected across four adult hospitals within the MLH system. We captured heart rate, SBP (via cuff if arterial not available), temperature, respiration rate and oxygen saturation based on availability of the data. In addition, the WBC count was collected from patients’ electronic health records. The onset of sepsis was not recorded in the dataset. Instead, we used the time that a patient fulfilled SIRS criteria to estimate the time of sepsis if it occurred. We excluded patients without at least 8 hours of continuous data, or who had ICD 10 codes in their discharge summary that indicated cardiovascular disease, including but not restricted to: congestive heart failure, arrhythmias, and myocardial infraction (see consort diagram in the Supplemental Figure 1). IRB approval was received for this retrospective observational study.

### 2.2 Features Extracted from Moving Time Windows

We allow the time-window to span two hours. Each minute, data from the moving time-window were used as a model input. These inputs included 38 parameters (basic statistics and signal information) describing the entire two-hour time interval in addition to 35 parameters (basic statistics) from the last 60 and 30 minutes of the time-window. These processed data were combined, making up a total of (38 + 35×2=)108 features, which served as the features of each two-hour time-window.

We analyzed data from two-hour moving time-windows for an overall observational period covering three and six hours and developed two machine learning models. As seen in Figure 1, the developed models would start producing results two hours into the observational period. Longer observational periods were considered, however, they resulted in poor data availability.

### 2.3 Random Forest Model

The random forest (RF) algorithm is a supervised classification algorithm that relies on the aggregate classification or the ‘majority vote’ of a series of decision trees, each built using random subsets of the dataset [26]. In general, RF algorithms outperform other tree-based classification algorithms such as decision tree learning [27] and tree bagging [28]. Random forest algorithms are generally robust against overfitting and are not overly sensitive to noisy data or a high ratio of parameters to observations [29]. In addition, RF algorithms can be used to identify and rank the most important features contributing to classification performance. Most importantly, compared to machine learning algorithms such as neural networks or other deep learning techniques, RF algorithms are known for their fast training and computation time for large datasets [30, 31], making them particularly appropriate for ‘online’ sepsis prediction work.

To objectively evaluate the models, 80% of the data were used for training the models and the remaining 20% were used for testing. In addition, the number of trees in the RF models, chosen based on the out-of-bag (OOB) error (26), was set to 700.

### 2.4 Performance Metrics

To evaluate the model performance, we used sensitivity, specificity, positive predictive value (PPV), accuracy, and F1 score of the developed RF algorithms when applied to the test sets. Sensitivity measures the proportion of correctly predicted sepsis patients from the sepsis group. Similarly, specificity measures the proportion of correctly predicted non-sepsis patients among the non-sepsis group. Positive predictive value measures the proportion of sepsis patients among those predicted as positives. Accuracy gives the overall proportion of correct predictions. Lastly, F1 score, which is another measure of a test accuracy, is the harmonic mean of sensitivity and PPV.

## 3. Results

Table 1 illustrates several descriptive statistics for the sepsis and non-sepsis patient subgroups for the 3-hour observational period. A *t*-test between the sepsis and non-sepsis subgroups indicate statistically significant differences across all parameters based on p-values.

Balanced training and test sets, with equal number of sepsis and non-sepsis patients, were generated to avoid favoring the more represented observations in the dataset. As discussed in Section 2.2, in this study, two observational periods of lengths three and six hours were used. For instance, for the sepsis case group, we selected a 3-hour observational period prior to sepsis onset, and in the control group, which consists of non-sepsis patients, we used a given 3-hour observational period. Hence, at the end of the observational period, half of the patients developed sepsis. A total of 377 and 293 sepsis patients were included for the 3- and 6-hour observational periods, respectively (see Supplemental Figure 1).

Figure 2 presents the F1 score, accuracy, PPV, sensitivity, and specificity of the RF algorithms when applied to the test set in an online fashion for 3- and 6-hour observational periods. As seen in Figure 2(a), for the 3-hour observational period, F1 score, accuracy, and sensitivity generally increase as a subset of patients start to develop sepsis while PPV and specificity are consistently high, ranging between 85%-94%. For instance, as seen in Figure 2(a), the algorithm developed using the 3-hour observational period can discriminate patients with accuracy 79% and F1 score 75% half an hour before the onset of sepsis. These percentages increase to 81% and 79%, respectively, 10 minutes before sepsis onset. Similarly, sensitivity equals 65% and 70% half an hour and 10 minutes before the onset of sepsis. Similarly, in Figure 2(b), for the 6-hour observational period, F1 score, accuracy, sensitivity, and PPV start to increase 1-1.5 hours before sepsis onset as a subset of patients start to exhibit physiomarkers while specificity is consistently high throughout the observational period. The algorithm can discriminate patients with accuracy 72% and F1 score 69% half an hour before the onset of sepsis. These percentages increase to 78% and 76%, respectively, 10 minutes before sepsis onset. Lastly, sensitivity, which is as low as 46% four hours before the onset of sepsis, increases to 63% and 71% half an hour and 10 minutes before the onset of sepsis. Note that the algorithms developed using 3- and 6-hour observational periods are similar in performance; however, the former is slightly better as it relies on a higher number of patients.

Figure 3 presents the probability of developing sepsis for an example sepsis patient as estimated by the model in an online fashion during a 3-hour observational period. As seen in the figure, the probability of developing sepsis is consistently low and then dramatically increases approximately 40 minutes before the sepsis onset. In this study, we predict sepsis to be present when the model-estimated probability of sepsis goes above 0.5. Hence, as seen in Figure 3, for this example patient the algorithm would predict sepsis 41 minutes before onset, allowing for directing the attention of the clinical staff to the patient to possibly prevent sepsis development and improve patient outcomes. In general, according to our retrospective analysis, when the developed RF model is able to predict sepsis, it does so on average (with a 95% confidence interval) 48.60 ± 4.20 minutes and 210.23 ± 17.83 minutes before the onset for the 3- and 6-hour observational periods, respectively.

**Figure 2.**
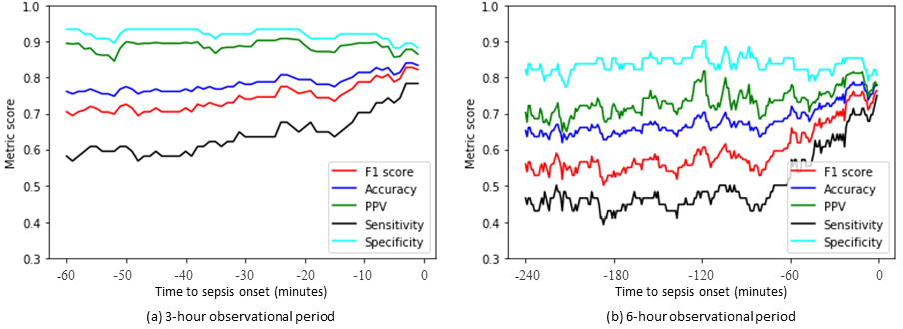
F1 score, accuracy, PPV, sensitivity, and specificity of the trained RF classifiers in discriminating sepsis and non-sepsis patients when used in an online fashion on separate test sets for 3- and 6-hour observational periods.

**Figure 3.**
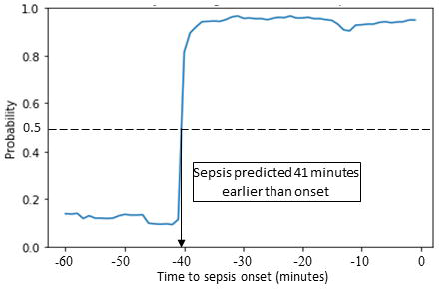
The model-estimated probability of developing sepsis for a given sepsis patient over the 3- hour observational period leading to sepsis onset

Figure 4 presents the aggregated mean and 95% confidence interval of the model-estimated probabilities for the test sets, in an online fashion during the 3-hour observational period, stratified across the sepsis and non-sepsis subgroups. As seen in Figure 4(a), the probability of being classified as an at-risk sepsis patient increases for the sepsis subgroup during the observational period as the time of sepsis onset approaches. In contrast, as illustrated in Figure 4(b), the probability of being classified as an at-risk sepsis patient for the non-sepsis subgroup stays relatively constant between 20% and 22% during the entire observational period. In addition, the confidence interval width decreases for the sepsis subgroup as the time to sepsis onset approaches and physiomarkers become more apparent, illustrating more confidence in the model estimated probabilities. In contrast, the confidence interval width for the non-sepsis subgroup slightly increases as the time to sepsis onset approaches.

**Figure 4.**
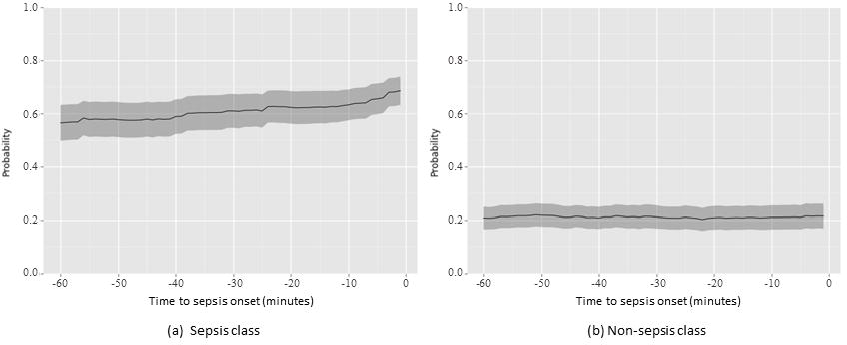
Mean and 95% confidence interval of the model-estimated probabilities for the test set stratified across the sepsis and non-sepsis subgroups for the 3-hour observational period.

## 4. Discussion

Continuous monitoring applied to the development of sepsis in hospitalized patients can greatly reduce the gap between biological onset and clinical recognition. In this study we demonstrate that physiomarkers exist prior to the onset of sepsis. Furthermore, the model appears to generate relatively few false positives, which would minimize alert fatigue. Specifically, the results indicate that the model is able to discriminate the case and control group with accuracy and F1 score of up to 79% and 75% half-hour before sepsis onset, and 81% and 79% ten minutes before sepsis onset. Note that in practice, sepsis onset itself may go unnoticed as it relies on continuously calculating SIRS criteria, which is not necessarily implemented in many early warning systems. Hence, if implemented in practice, the developed automated algorithms can result in improved outcomes beyond the presented results.

The feature extraction used in this study is novel and inspired by the body’s temporal response to infection. Specifically, we extracted features from high-frequency data streams during relatively large time-window, i.e., two hours, to establish a proper baseline. However, we also combined these extracted features with those obtained from shorter time intervals, i.e., the last 60 and 30 minutes within the time-window, to emphasize the latest physiological changes. The impact and benefit of this feature extraction approach is also evident based on the most significant features contributing to discriminating between sepsis and non-sepsis patients. Our analysis shows that the top physiological features in both models developed using the 3- and 6-hour observational periods are drawn from a mixture of these different interval sizes. Specifically, for the 3-hour observational period, the top physiological features include the maximum respiratory rate from each of the 2, 1, and 0.5-hour interval windows, as well as the variance of SBP and mean heart rate for the 0.5-hour interval window. Similarly, for the 6-hour observational period, the top physiological features include the maximum respiratory rate from each of the 2, 1, and 0.5-hour interval windows and the mean and maximum heart rate from the 0.5-hour interval window (see Supplemental Table 3).

Furthermore, a unique advantage of our models is the minimal set of required variables, which only include four routinely collected physiological data (heart rate, respiratory rate, SBP, and temperature) and basic laboratory data, i.e., WBC count. In other words, the models do not require additional laboratory data such as creatinine level, or subjective clinical metrics such as Glasgow Coma Scale that are usually required to meet more complicated severity of illness criteria. The use of a minimal set of variables in our model ultimately makes it easier to implement at the bed-side and without significant manual data entry.

The developed models show promise in the early prediction of sepsis, providing an opportunity for directing early intervention efforts to prevent/treat sepsis cases. However, more study is needed as our work has limitations. In this study, while four hospitals were involved in the data collection, all patients were selected from a single hospital system in the Mid-South USA. Moreover, all patients were admitted to the emergency department, ICU, surgical wards, or other complex care units. Furthermore, the models were developed using the data from patients for whom SIRS criteria could be calculated. We also identified suspicion of infection by referring to the presence of ICD 10 discharge codes for sepsis and severe sepsis. We recognize that it is possible that some patients who developed sepsis during their stay were not coded. Also, the current developed models require WBC count to discriminate patients; however, future models can be developed without reliance on WBC count. In addition, we use continuous minute-by-minute physiological data streams. Such facilities may not be available at all hospital settings; hence our work only applies to facilities where streaming data is available. Finally, the models were developed using a retrospective dataset, hence prospective application would reveal significant information about the practical utility and effectiveness.

In conclusion, we illustrate the performance of a state of the art machine learning technique applied to data captured at the bedside with a short collection interval. We develop RF models for various time periods before sepsis onset, in order to discriminate between sepsis and non-sepsis patients. These results demonstrate that salient physiomarkers identified in continuous physiological data streams can add significant value to decision making at the bedside by predicting patients at the highest risk for developing sepsis.

## Acknowledgements

We would like to acknowledge the efforts of Michael Younker, Brian Williams, Don MacMillan for their work in preparing and providing key data elements that were used in this paper

